# Photostimulation Improves Maturation of Human Photoreceptors

**DOI:** 10.1101/2025.07.17.665282

**Authors:** Canan Celiker, Eva Hruba, Konstantina Kyriakou, Birthe Dorgau, Francisco Molina Gambin, Kamila Weissova, Lucie Englmaier, Nadezda Vaskovicova, Michael A. Savage, Gerrit Hilgen, Evelyne Sernagor, Alejandro Garanto, Majlinda Lako, Tomas Barta

## Abstract

The human retina contains photoreceptor cells that detect light and enable vision. The development of these cells involves a tightly regulated cascade of structural and molecular events, and their dysfunction leads to irreversible blindness in many retinal diseases. Human retinal organoids derived from stem cells have become powerful tools to model retinal development and disease, but they often remain immature and lack key features required for full function. Light is not only the sensory target of photoreceptors but also an important developmental signal *in vivo*. However, light has rarely been used as a deliberate stimulus during *in vitro* differentiation. Here we show that exposing retinal organoids to rhythmic light flicker at a specific frequency enhances photoreceptor maturation across multiple levels. This stimulation improves the development of outer segments, accelerates the transcriptional transition from precursor to mature photoreceptors, and strengthens functional connectivity with downstream neurons. These findings identify patterned light as a potent and physiologically relevant signal for driving retinal development *in vitro*. This approach represents a non-invasive and easily scalable method for improving the quality of retinal organoids, with implications for disease modelling, drug discovery and the preparation of photoreceptors for cell-based therapies.

## Main

The human retina is a complex neural tissue that captures light and initiates visual processing through the action of photoreceptor cells—rods and cones—whose development requires precise orchestration of gene expression, polarity, morphogenesis, and synaptogenesis. Retinal organoids derived from human pluripotent stem cells have emerged as powerful models to study these processes *in vitro*, yet they often exhibit incomplete maturation, particularly in forming fully developed outer segments and establishing functional synapses ^1,2^. This developmental arrest is especially pronounced in long-term cultures and remains a bottleneck for both fundamental and translational applications. These limitations have far-reaching consequences. In disease modelling, immature photoreceptors may fail to replicate key pathological features of inherited retinal disorders, limiting the utility of organoids for studying disease progression or testing therapeutic interventions. In drug discovery, insufficient maturation hampers the ability to assess compound efficacy on functional phototransduction or synaptic signalling. Critically, in the context of photoreceptor transplantation—a promising strategy for restoring vision in degenerative retinal diseases— the ability of donor cells to survive, integrate, and contribute to visual function largely depends on the maturity of cells and synaptic machinery^3,4^. Therefore, strategies that promote the full maturation of photoreceptors in retinal organoids are urgently needed to improve the fidelity, functionality, and clinical relevance of this model system.

While advances in organoid engineering have shown that physical and sensory cues—such as mechanical strain, electrical stimulation, or circadian entrainment—can accelerate the maturation of other stem cell-derived tissues (e.g., cardiomyocytes, neurons, β-cells)^5–10^, the role of light as a developmental stimulus on the retina has been largely overlooked. This is surprising given that light is not only the physiological target of photoreceptors but also an essential instructive signal during retinal development *in vivo*^11–15^.

Here, we address this fundamental and previously untested question: Can light—delivered in a controlled, rhythmic pattern—promote the maturation and function of human photoreceptors in retinal organoids? By applying 40 Hz flickering light in a 12-hour light / 12-hour dark cycle, we demonstrate that photostimulation markedly enhances photoreceptor maturation at structural, transcriptomic, synaptic, and electrophysiological levels. In doing so, we identify light not only as a sensory end-point but as a powerful developmental signal for guiding human retinal differentiation.

## Results

### Light Stimulation Improves Outer Segments Formation

Light is a fundamental driver of retinal development^12–16^, and we hypothesized that controlled stimulation using light could enhance the differentiation and maturation of photoreceptor cells in human retinal organoids. Our proposed model suggests that such stimulation improves the structural and functional properties of the retina, with optimal conditions promoting the formation of well-developed outer segments and enhancing the expression of photoreceptor-specific markers (**Fig. 1a**).

**Figure 1.**
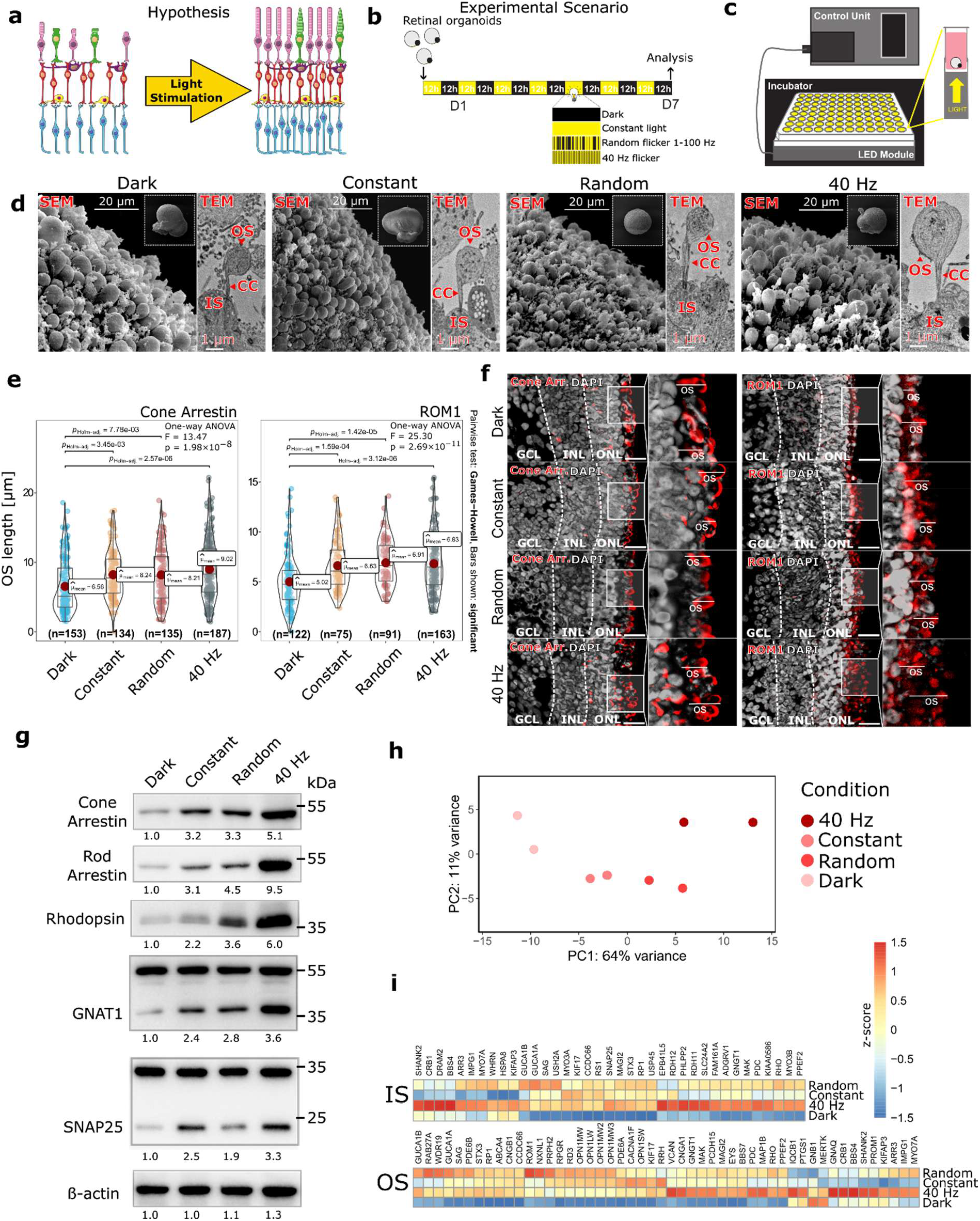
Light stimulation enhances retinal differentiation and outer segment maturation. **(a)** Schematic representation of the general hypothesis. **(b)** Experimental conditions used for light stimulation. Human retinal organoids were subjected to four conditions: darkness, constant light, random-frequency flickering light, and 40 Hz flickering light. **(c)** Light stimulation device (CellLighteR) used for delivering controlled light exposure to human retinal organoids. **(d)** Scanning electron microscopy (SEM) and transmission electron microscopy (TEM) images of retinal organoids demonstrating the impact of light stimulation on outer segment (OS) morphology. Dark-cultured organoids exhibit disorganized, short OS, while all light conditions promote elongation, with 40 Hz flickering light leading to the most mature and structured OS. Scale bars: SEM = 20 μm, TEM = 1 μm. IS = inner segment, CC = connective cilium. **(e)** Quantification of OS length using Cone Arrestin and ROM1 immunostaining. Statictics – one-way ANOVA, post hoc pair-wise Games-Howell. **(f)** Representative immunostaining images of Cone Arrestin and ROM1, corresponding to the quantification in (e). Scale bar = 20 μm. ONL = outer nuclear layer, INL = inner nuclear layer, GCL = ganglion cell layer. **(g)** Expression of Cone Arrestin, Rod Arrestin, Rhodopsin, GNAT1, and SNAP25 in retinal organoids in all tested light stimulation regimens, as determined using Western blot analysis. Numbers below show mean expression fold change, relative to dark condition. **(h)** Principal component analysis (PCA) plot showing the clustering of transcriptomic profiles of light stimulated retinal organoids. **(i)** Heatmap shows expression of major OS and IS-specific genes in retinal organoids upon different light stimulation regimes.

To test this hypothesis, we subjected human retinal organoids (**Fig. S1a–d**) to four different conditions: complete darkness, constant light, random-frequency flickering light (1–100 Hz), and flickering light at 40 Hz. We selected this gamma-band oscillation at 40 Hz because it represents a fundamental feature of neuronal network dynamics, known to facilitate sensory processing, synaptic plasticity, and circuit maturation^17–19^. All experiments were conducted using organoids between day (D) 150 and D180 of differentiation, a developmental window during which post-mitotic CRX+ photoreceptors are present and both inner and outer segment structures begin to form. Notably, retinal organoids become transcriptionally and electrophysiologically responsive to light between D120 and D150^20,21^, aligning with the onset of photoreceptor maturation. All conditions were applied in a 12-hour light / 12-hour dark cycle for 7 consecutive days to mimic natural day/night rhythms, and organoids were harvested at the end of the dark phase to minimize acute light-induced effects (**Fig. 1b**). All experiments were performed using our custom-built Cell LighteR device^20^, designed to deliver precise and controlled light stimulation to organoid cultures (**Fig. 1c**).

To assess the impact of light stimulation on photoreceptor maturation, we used outer segment (OS) length and structure as a rapid and reliable readout, analysing their morphology with scanning electron microscopy (SEM) and transmission electron microscopy (TEM) (**Fig. 1d**). Dark-cultured organoids displayed short OS-like structures, while all light-treated groups exhibited significant improvements in OS length and organisation. Notably, the 40 Hz flickering condition resulted in the most well-defined and elongated OS structures (**Fig. 1d**). To quantify these changes, we measured OS length using ROM1 and Cone Arrestin (gene *ARR3*) immunostaining (**Fig. 1e, f**). Compared to the dark condition, all light-exposed groups showed significantly elongated OS, with the most pronounced effect observed under 40 Hz flickering light. Immunostaining for critical photoreceptor proteins further confirmed the differentiation-enhancing effect of light. Compared to dark-cultured organoids, all light conditions resulted in increased expression of key photoreceptor markers, including rhodopsin (RHO), GNAT1, PRPH2, and PDE6A, as well as inner segment (IS) mitochondrial marker TOMM20 (**Fig. S1e**). The expression levels of Cone Arrestin, Rod Arrestin (gene *SAG*), Rhodopsin, GNAT1, and the synaptic marker SNAP25 were all elevated in light-treated conditions compared to the dark group (**Fig. 1g, S2**). Among all conditions, 40 Hz flickering light led to the highest protein expression levels.

To obtain a global overview of transcriptomic changes, we conducted bulk RNA sequencing. Principal component analysis (PCA) of the transcriptome clearly segregated samples by stimulation condition (**Fig. 1h**). The 40 Hz light group formed a distinct cluster, indicating that this condition induces the most pronounced and consistent transcriptional changes among all tested paradigms. Expression changes of genes associated with photoreceptor OS and photoreceptor inner segment (IS) structures, as defined by Gene Ontology annotations (https://www.ebi.ac.uk/QuickGO/), were visualized using a heatmap **(Fig. 1i**). The majority of these genes were significantly upregulated under light stimulation, with the strongest induction observed in the 40 Hz flickering condition.

Importantly, all experiments were conducted in phenol red-free medium, as we observed that the presence of phenol red markedly interfered with the effects of light. When experiments were carried out in medium containing phenol red, no significant differences in gene expression were detected across conditions. Specifically, the expression levels of key photoreceptor genes— *RHODOPSIN, OPN1SW*, and *SAG*—remained uniformly low, indicating that light failed to elicit the expected transcriptional upregulation in the presence of phenol red (**Fig. S3**).

### Flickering Light at 40 Hz Induces Cell-Type Specific Responses in Retinal Organoids

Among all tested conditions, 40 Hz flickering light consistently demonstrated the most pronounced effects on photoreceptors (**Fig. 1**). Therefore, we selected the 40 Hz condition as the most effective stimulation paradigm and focused subsequent analyses on uncovering its cell-type-specific impact.

We performed single-cell RNA sequencing (scRNA-seq) on human retinal organoids subjected to either dark or 40 Hz flickering light (further referred as photostimulation) in a 12-hour light / 12-hour dark cycle for 7 days (**Fig. 2a**). Using the human retina atlas gene annotations^22^, we identified 13 distinct retinal cell types, including rod photoreceptors, cone photoreceptors, rod and cone precursors, horizontal cells, amacrine cells, retinal ganglion cells, (ON and OFF) bipolar cells, Müller glia, astrocytes, retinal pigment epithelium (RPE), and retinal precursors (**Fig. 2b,c, Table S1**). Interestingly, photostimulation led to an increase in rod and cone photoreceptors at the expense of their respective precursor populations, indicating accelerated differentiation under flickering light conditions (**Fig. 2d, 3c**). In contrast, the proportion of Müller glial cells was reduced in photostimulated organoids, potentially indicating decreased gliosis or a shift in cellular composition favouring neuronal differentiation (**Fig. 2d**). To identify genes differentially expressed between dark and photostimulated conditions, we first analysed global transcriptomic changes across all cells (**Fig. 2e, f**). Key photoreceptor-associated genes were upregulated in response to flickering light exposure, including *PDE6G, SAG, PDE6A, PRPH2, ARR3, TTR, RHO, ESRRB*, and *GUCA1B*. These genes are crucial for phototransduction, outer segment development, and photoreceptor maintenance, suggesting that photostimulation enhances photoreceptor function and maturation. Conversely, several genes characteristic of Müller glia, including *VIM* (vimentin) and *CRABP1*, were downregulated in photostimulated condition, potentially corroborating the observed reduction in the Müller glial population. Surprisingly, we did not observe profound upregulation of genes associated with oxidative stress or apoptosis upon photostimulation (**Fig. S4a, b**). Gene ontology (GO) analysis of all significantly differentially expressed genes (p < 0.05) revealed strong enrichment for sensory perception of light, visual perception, detection of light stimulus, phototransduction, and photoreceptor cell maintenance, supporting the role of flickering light in promoting photoreceptor functionality (**Fig. 2g**). Additionally, GO analysis for cellular components showed that differentially expressed genes were significantly associated with photoreceptor outer segments, photoreceptor cilia, photoreceptor disc membranes, and ciliary membranes, highlighting the structural and functional improvements in photoreceptors following light stimulation (**Fig. 2h**).

**Figure 2.**
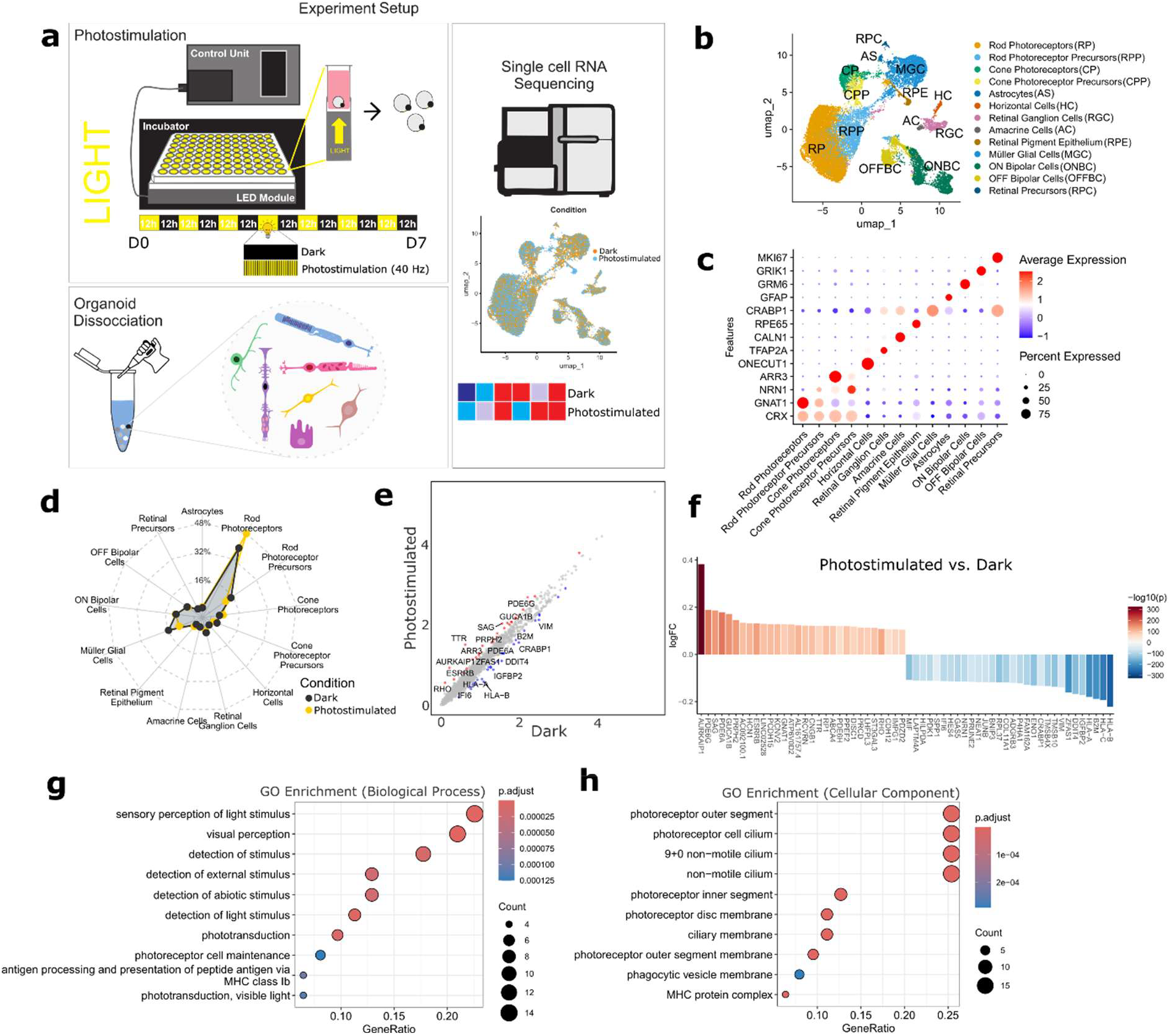
Single-Cell RNA sequencing reveals specific responses to photostimulation. **(a)** Experimental setup for scRNA-seq analysis. Retinal organoids were cultured under dark or 40 Hz flickering light (Photostimulation) conditions for 7 days (12-hour light / 12-hour dark cycle), followed by dissociation and single-cell RNA sequencing. **(b)** UMAP projection of retinal organoid cells, identifying 13 distinct retinal cell types. **(c)** Dot plot showing marker gene expression for cell type validation. **(d)** Cell-type abundance analysis comparing dark vs. photostimulated. **(e)** Differential gene expression analysis comparing dark vs. photostimulated. **(f)** Waterfall plot displaying the top 30 differentially expressed genes in each condition. **(g)** Gene Ontology (GO) enrichment analysis for biological processes. **(h)** GO enrichment analysis for cellular components.

### Photostimulation improves maturation of photoreceptors

To investigate the effect of photostimulation on photoreceptors, we focused our single-cell transcriptomic analysis on rod and cone lineages. Sub-clustering of the photoreceptor population revealed four distinct cell types corresponding to rod photoreceptors, cone photoreceptors, and their respective precursors (**Fig. 3a**). The overall transcriptional landscape showed clear separation between mature and immature photoreceptors, enabling a detailed examination of differentiation dynamics. Pseudotime trajectory analysis projected onto the UMAP embedding revealed a continuous transition from photoreceptor precursors to mature cells (**Fig. 3b**). Early pseudotime values were associated with precursor clusters, while high pseudotime scores marked terminally differentiated photoreceptors. Notably, cells from the photostimulated group were enriched in late pseudotime regions, indicating accelerated maturation. A quantitative comparison of cell counts in the clusters under dark and stimulated conditions further supported this shift (**Fig. 3c**). Numbers of rod and cone photoreceptors were profoundly increased in photostimulated samples, whereas precursor populations were overrepresented under dark conditions. This redistribution suggests that photostimulation promotes photoreceptor terminal differentiation.

**Figure 3.**
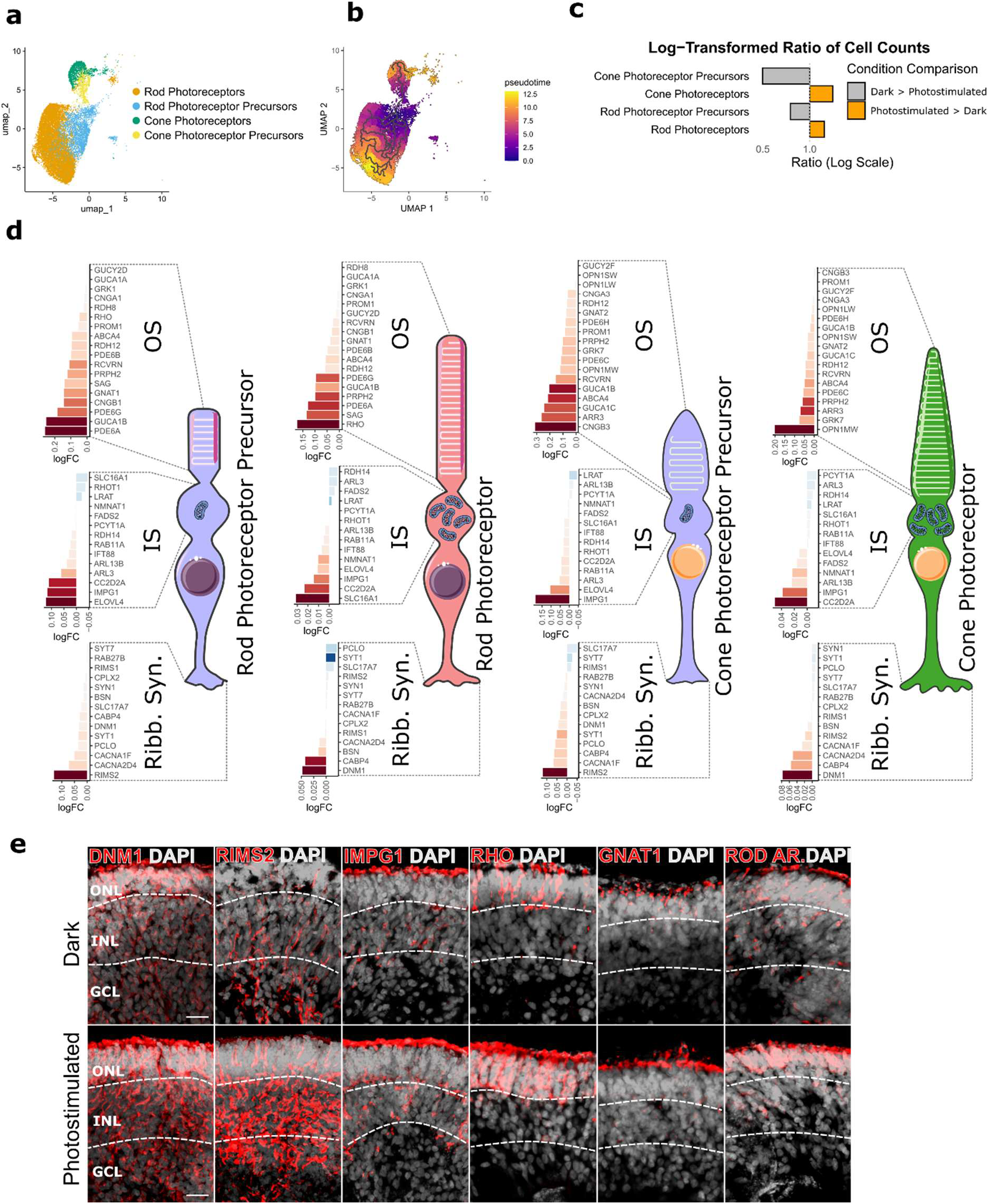
Photostimulation promotes maturation of rod and cone photoreceptors. **(a)** UMAP embedding of sub-clustered photoreceptors, identifying four transcriptionally distinct populations: rod photoreceptors, cone photoreceptors, rod photoreceptor precursors, and cone photoreceptor precursors. **(b)** Pseudotime trajectory analysis overlaid onto the UMAP space reveals a continuous developmental progression from precursor to mature photoreceptor states. **(c)** Log-transformed number of cells in each photoreceptor cluster under dark and photostimulated conditions, showing enrichment of photoreceptors in the photostimulated group and accumulation of precursors in the dark condition. **(d)** Schematic summary of gene expression changes in each photoreceptor population. For each subtype, three waterfall plots depict photostimulation-induced transcriptional changes in major genes associated with outer segments (OS), inner segments (IS), and ribbon synapses. **(e)** Immunostaining for DNM1, RIMS2, IMPG1, RHODOPSIN (RHO), GNAT1, and ROD ARRESTIN (ROD AR.) in retinal organoid sections. ONL = outer nuclear layer, INL = inner nuclear layer, GCL = ganglion cell layer. Scale bar = 20 μm.

To examine subtype-specific responses to light in more detail, we generated schematic representations of each photoreceptor category, overlaying gene expression changes induced by photostimulation (**Fig. 3d**). Waterfall plots illustrated consistent upregulation of major genes associated with OS, IS, and ribbon synapse development across all cell subtypes. These included functional markers such as *RHO, SAG, GNAT1, ARR3, PDE6A, IMPG1, RIMS2*, and *DNM1*, highlighting improved structural and synaptic maturation in response to stimulation. To validate these transcriptomic findings at the protein level, we performed immunohistochemistry for RHODOPSIN, GNAT1, ROD ARRESTIN, DNM1, RIMS2, and IMPG1—key proteins involved in OS, synaptic vesicle recycling, ribbon synapse organization, and interphotoreceptor matrix composition, respectively. All markers showed visibly increased expression and broader distribution in photostimulated retinal organoids compared to dark controls, with strongest signal observed in layers corresponding to the outer nuclear layer and synaptic regions (**Fig. 3e**). These results support the conclusion that photostimulation enhances the structural and functional maturation of photoreceptors at both transcriptional and protein levels.

To validate the inferred maturation trajectory, we assessed the spatial expression of canonical photoreceptor markers across the UMAP embedding. CRX, a key regulator of photoreceptor fate, was broadly expressed across all photoreceptor populations (**Fig. S5a**). In contrast, *SAG* (rods) and *ARR3* (cones) were restricted to terminally differentiated cells, while *PTCHD4, FOXD1*, and *BHLHE40* were confined to precursor clusters (**Fig. S5a–b**). Differential expression analysis between precursors and photoreceptors confirmed the upregulation of phototransduction, synaptic, and structural gene networks in differentiated cells (**Fig. S5b**).

To evaluate how stimulation modulates gene expression within each photoreceptor type, we compared transcriptomic profiles of photostimulated and dark-cultured organoids. UMAP plots with overlaid condition labels showed an enrichment of photostimulated cells in mature photoreceptor clusters and a reciprocal enrichment of dark-cultured cells in precursor clusters (**Fig. S6a–b**). Expression of key functional genes—including *SAG* and *ARR3* — was elevated in photostimulated cells, whereas precursor-associated genes (*PTCHD4, FOXD1, BHLHE40*) were diminished (**Fig. S6c-f**).

Collectively, these data demonstrate that photostimulation promotes the terminal differentiation of rod and cone photoreceptors by accelerating pseudotime progression, modulating transcriptional signatures, and increasing the proportion of mature cell types. This maturation-enhancing effect is reflected in both global cluster distribution and the activation of key photoreceptor-specific gene networks.

### Photostimulation improves electrophysiological properties

Having observed transcriptional and structural maturation of photoreceptors in response to photostimulation, we next assessed whether these changes translated into functional improvements at the synaptic and electrophysiological level. We first examined the expression of genes associated with synaptic transmission, with particular focus on components of the photoreceptor ribbon synapse. Dot plots comparing dark and photostimulated conditions revealed profound upregulation of key synaptic genes, including *STX3, BSN, SYP*, and *DLG4 (PSD95)*, in both rods and cones following stimulation (**Fig. 4a**). These genes are essential for vesicle docking, neurotransmitter release, and synaptic scaffold formation, suggesting that stimulation enhances synaptogenesis and synaptic maturity in photoreceptors. To validate these transcriptional changes at the protein level, we performed immunostaining for the same synaptic markers in retinal organoids cultured under either dark or photostimulated conditions. All four proteins— STX3, BSN, SYP, and PSD95—showed stronger and more widespread expression between the outer nuclear layer (ONL) and the inner nuclear layer (INL) (presumptive outer plexiform layer) of photostimulated organoids compared to dark controls (**Fig. 4b**).

**Figure 4.**
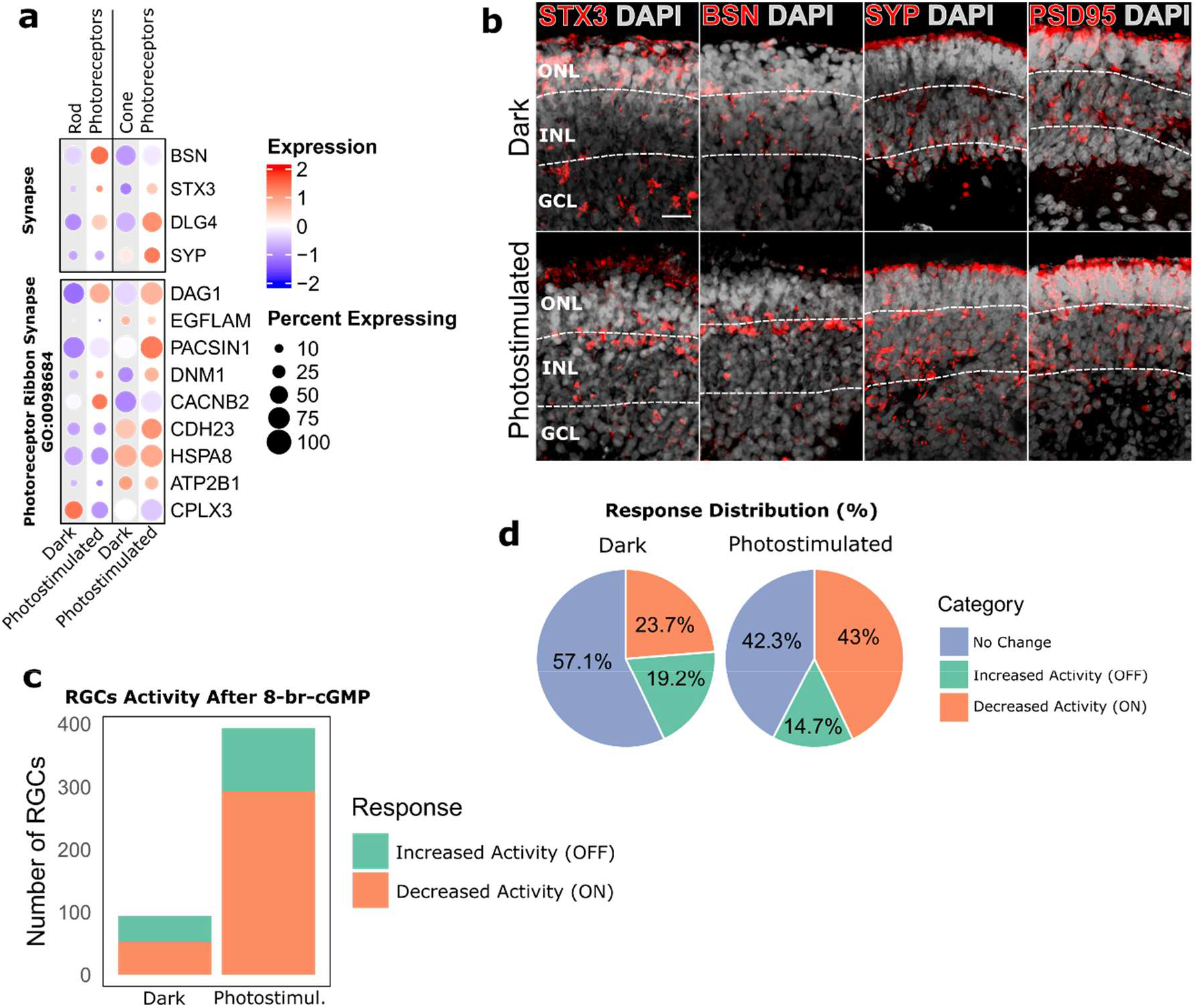
Photostimulation enhances synaptic maturation and retinal ganglion cell (RGC) responsiveness. **(a)** Dot plot showing differential expression of synaptic genes, including ribbon synapse components, in rod and cone photoreceptors under dark and photostimulated conditions. **(b)** Immunostaining for synaptic markers STX3, BSN, SYP, and PSD95 in retinal organoid sections. All markers exhibit increased signal intensity and broader distribution in photostimulated organoids, indicating enhanced synaptic development. ONL = outer nuclear layer, INL = inner nuclear layer, GCL = ganglion cell layer. Scale bar = 20 μm. **(c)** Quantification of RGCs recorded using high-density multi-electrode array (MEA-HD). **(d)** Distribution of RGC response types under dark and photostimulated conditions.

To determine whether the improved cellular and synaptic maturation observed with photostimulation translates into enhanced retinal output, we performed high-density multi-electrode array (MEA-HD) recordings on retinal organoids maintained under dark or photostimulated conditions. To ascertain that the recorded responses originated from photoreceptors and not from intrinsically photosensitive retinal ganglion cells (ipRGCs), we applied 8-bromoguanosine 3′,5′-cyclic monophosphate (8-br-cGMP), a membrane-permeable analogue of cGMP, into the recording chamber. cGMP is a key second messenger in the phototransduction cascade, gating Na+-permeable channels in photoreceptor outer segments. Puffing 8-br-cGMP mimics the dark current by depolarizing photoreceptors through Na+ influx, thereby initiating neurotransmitter release to downstream neurons^21^. This approach allows for the selective activation of photoreceptor-driven circuitry independent of actual light exposure. RGCs that responded to 8-br-cGMP with a ≥50% decrease in spiking activity were classified as presumptive ON cells, consistent with their expected hyperpolarization upon photoreceptor activation. Conversely, RGCs that exhibited a ≥50% increase in firing rate were categorized as OFF cells (**Fig. S7**). MEA recordings revealed a greater number of responsive RGCs in the photostimulated group compared to dark controls, with both ON-and OFF-type responses more prevalent (**Fig. 4c**). This shift was further reflected in the distribution of response types, with photostimulation yielding a higher proportion of ON RGCs—suggesting enhanced functional integration of photoreceptors within the retinal circuitry (**Fig. 4d**). These results support the conclusion that photostimulation enhances photoreceptor differentiation and improves the functional connectivity and responsiveness of the retina.

### Photostimulation engages the circadian machinery in retinal organoids

Our photostimulation paradigm was applied using a 12-hour light / 12-hour dark cycle, a strategy designed not only to mimic natural diurnal rhythms but also to engage the intrinsic circadian machinery of the retina. *In vivo*, photoreceptors are tightly regulated by the circadian clock, which coordinates daily fluctuations in gene expression, outer segment turnover, and synaptic transmission^23,24^. We therefore hypothesized that retinal organoids exposed to rhythmic light might entrain endogenous circadian oscillators, further supporting their maturation and physiological fidelity.

To investigate whether human retinal organoids possess a functional molecular clock that can respond to external cues, we employed a *BMAL1*-driven luciferase reporter system. Human iPSCs were transduced with a lentiviral construct encoding the *BMAL1*::luciferase cassette, and luciferase activity was monitored in real time using a custom-adapted version of the LuminoCell^25^ – a platform capable of measuring luciferase activity in individual organoids in a 96-well plate format (**Fig. 5a**). We first examined early-stage retinal organoids (day 20) to assess whether the core circadian transcriptional machinery was active or inducible. While untreated controls showed no rhythmicity, organoids synchronized with dexamethasone (DEX) displayed robust circadian oscillations in *BMAL1*-driven bioluminescence (**Fig. 5b**), indicating that a functional molecular oscillator is already present at this early stage.

**Figure 5.**
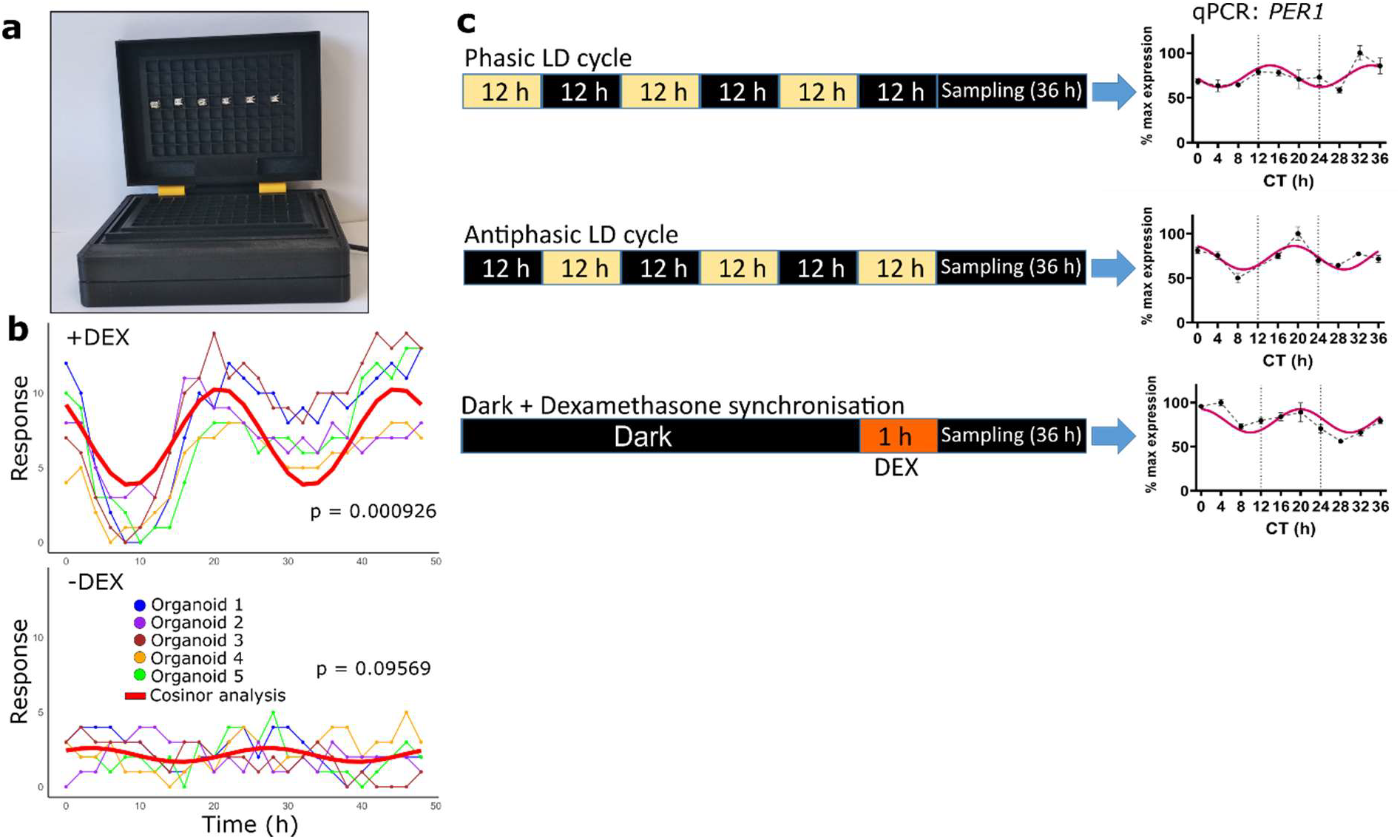
Retinal organoids possess functional circadian machinery responsive to photic cues. **(a)** Photograph of a custom-modified LuminoCell device adapted for bioluminescence monitoring of individual retinal organoids in a 96-well format. The device continuously records luciferase activity driven by a BMAL1 promoter. **(b)** Bioluminescence traces from five day-20 retinal organoids expressing a BMAL1::luciferase reporter, either synchronized with dexamethasone (+DEX) or left unsynchronized (−DEX). Rhythmic oscillations are observed only in the synchronized group. Cosinor analysis confirms statistically significant rhythmicity in the +DEX condition. **(c)** qPCR analysis of PER1 expression in day-180 retinal organoids following circadian synchronization. Organoids were exposed to either phasic or antiphasic 12 h light / 12 h dark cycles for 48 hours, followed by monitoring under constant darkness. Dexamethasone-treated organoids served as a positive control. Both light-synchronized groups exhibited distinct oscillatory patterns of PER1 expression, indicating successful circadian entrainment by light. served as positive controls (**Fig. 5c**). These findings demonstrate that light is sufficient to entrain circadian gene expression in retinal organoids and that phase relationships are preserved across experimental conditions.

To determine whether light could act as an entraining cue in more mature organoids, we exposed day 180 retinal organoids to a 72-hour light-dark cycle—sufficient to synchronize circadian rhythms^8,9^—followed by 36 hours of sampling under constant darkness at 4-hour intervals. Using qPCR, we tracked the expression of *PER1*, a canonical circadian clock gene, in organoids that had been synchronized either in phase or in antiphase (**Fig. 5c**). In both cases, we observed clear oscillations in *PER1* transcript levels, while dexamethasone-treated samples

Together, these results confirm that the circadian machinery in human retinal organoids is functional and light responsive. The use of a 12-hour light/dark stimulation protocol therefore serves a dual role—both as a physiological entrainment signal and as a maturation-enhancing cue—supporting the idea that rhythmic environmental inputs can drive developmental and functional refinement in stem cell-derived neural tissues.

## Discussion

In this study, we demonstrate that temporally structured light, specifically 40 Hz flickering light, significantly enhances the differentiation and maturation of human photoreceptors in retinal organoids. Our findings reveal that light is not only the physiological target of photoreceptors but also a potent environmental cue capable of guiding their development. Among several stimulation paradigms tested—including constant illumination and randomly flickering light— gamma-frequency flicker (40 Hz) emerged as uniquely effective in promoting outer segment elongation, increasing expression of photoreceptor-specific genes, and improving electrophysiological properties. These effects were accompanied by increased synaptic protein expression and enhanced responsiveness of downstream retinal ganglion cells, indicating a systems-level improvement in organoid functionality.

The concept of using environmental stimulation to promote cell differentiation is well established in other organoid and stem cell systems. Mechanical stretch and electrical stimulation enhance sarcomere formation and alignment in cardiomyocytes^5,7^, electrical pulses increase neuronal excitability and accelerate the development of synaptic networks^6^, and circadian metabolic entrainment through time-restricted feeding supports the functional maturation of β-like cells in pancreatic islets^9^. These approaches apply physical or rhythmic inputs that cells would normally encounter *in vivo*. Light, uniquely suited to the visual system, has been largely overlooked in this context. Although previous work has shown that early light exposure can influence photoreceptor survival, and connectivity in animal models^14,15^, this is, to our knowledge, the first systematic study showing that rhythmic light stimulation—specifically in the gamma frequency range—can drive terminal photoreceptor differentiation and enhance circuit-level responsiveness in human retinal organoids.

Our findings suggest that 40 Hz flickering light is particularly effective due to its resonance with intrinsic neural oscillations. Gamma-band oscillations (~30–80 Hz) are fundamental features of neuronal activity in the central nervous system and play critical roles in sensory processing, attention, and circuit maturation^18,26^. In the developing visual system, spontaneous and evoked retinal waves help pattern connectivity before photoreceptors become light-responsive^27,28^. Flicker at 40 Hz, which entrains gamma oscillations, has been shown to enhance synaptic plasticity, improve memory, and reduce amyloid and tau pathology in mouse models of Alzheimer’s disease^19,29,30^. Our data suggest that similar entrainment may also apply to immature retinal circuits, potentially activating downstream signalling or activity-dependent transcriptional programs necessary for photoreceptor maturation. Interestingly, in a complementary study by van Oosten et al. (submitted to BiorXiv at the same day as this work), they demonstrate that non-flickering, long-term photostimulation also promotes photoreceptor maturation in retinal organoids. While their study focuses on constant light exposure from day 70 of organoid differentiation process, the convergence of our results underscores the broader principle that light itself—whether rhythmic or continuous—acts as a potent developmental cue for human photoreceptors. Together, these findings highlight the versatility of photostimulation strategies and suggest that multiple temporal patterns of light may be harnessed to support retinal development, depending on the specific maturation stage or cellular context.

Single-cell transcriptomic analysis revealed that photostimulation alters cell-type proportions by increasing mature rod and cone photoreceptors while reducing precursor populations. This redistribution along the pseudotime trajectory indicates that light accelerates terminal differentiation. Notably, we also observed a reduction in Müller glia under photostimulated conditions, which could reflect reduced gliosis or a shift in lineage allocation—a phenomenon observed in other models where enhanced neuronal maturation reduces glial expansion^10^. Furthermore, transcriptomic signatures of phototransduction, OS structure, and ribbon synapses were all enriched following photostimulation, indicating comprehensive effects on photoreceptor architecture and function.

At the systems level, we found that light-exposed organoids exhibited enhanced expression of synaptic markers, including STX3, BSN, and PSD95, and generated more robust responses in downstream retinal ganglion cells (RGCs) when pharmacologically stimulated. These data are consistent with improved circuit connectivity and synaptic transmission. Similar enhancements in electrophysiological properties have been observed in brain organoids exposed to electrical stimulation^31^, and in cardiomyocytes following rhythmic pacing^5^. Our data suggests that a similar principle applies in the visual system demonstrating the utility of temporally structured stimuli for accelerating *in vitro* maturation.

One of the most promising therapeutic strategies for retinal degenerative diseases is the transplantation of photoreceptor precursors or mature photoreceptors derived from stem cells. However, despite decades of progress, the efficiency of photoreceptor transplantation remains limited. Challenges include poor cell survival, inadequate integration into the host outer nuclear layer, and limited functional synapse formation with downstream neurons^3,4,32,33^. A critical bottleneck is the maturity of the donor photoreceptors: cells that lack fully developed outer segments, proper polarity, or synaptic machinery are unlikely to survive transplantation or restore vision effectively. In addition to pre-conditioning donor cells *in vitro*, a key future direction will be to explore whether *in vivo* photostimulation—particularly with structured flickering light—could further enhance integration and functionality after transplantation. Our data indicate that 40 Hz flickering light promotes photoreceptor maturation and circuit engagement in organoid systems. It is conceivable that a similar stimulus, applied post-transplantation, could entrain host and donor retinal activity, promote synaptogenesis, and strengthen functional coupling with bipolar or ganglion cells. Such an approach would parallel therapeutic sensory entrainment strategies used in other areas of the nervous system, including auditory and cortical rehabilitation. The feasibility of delivering flickering light non-invasively and in a frequency-specific manner makes it an attractive candidate for adjunctive therapy in cell-based retinal repair. Preclinical models will be essential to determine the safety, optimal parameters, and efficacy of such interventions. Our findings suggest that pre-conditioning photoreceptor cells with structured photostimulation—particularly 40 Hz flickering light—can promote key aspects of photoreceptor maturation, including outer segment formation, expression of phototransduction machinery, and functional connectivity. This strategy may substantially enhance the readiness of stem cell-derived photoreceptors for transplantation by bringing them closer to a mature, transplantation-competent state.

Another promising avenue for improving photoreceptor maturation involves the modulation of culture media composition. Previous work by West and colleagues^34^ demonstrated that supplementation with specific lipids enhances OS formation and disk organization in developing photoreceptors. Given that OS morphogenesis depends on high membrane turnover and tightly regulated lipid dynamics, integrating lipid supplementation into photostimulation protocols could further synergize structural maturation. Combining temporal photostimulation with metabolic support through targeted lipid additives may promote both the elongation and lamellar organization of OS membranes, yielding photoreceptors with more native-like architecture. Exploring such multimodal approaches will be essential for optimizing the functional competence of retinal organoids and transplant-ready cells.

It is also important to highlight the technical simplicity and physiological relevance of the photostimulation paradigm. Unlike genetic modifications or pharmacological agents, patterned light is non-invasive, easily tuneable, and directly interpretable by the photoreceptor machinery. Importantly, we found that the presence of phenol red in the medium abolished the stimulatory effects of light—a finding consistent with reports that phenol red can act as a weak estrogen mimic or ROS scavenger ^35,36^.

While the effects of 40 Hz stimulation were robust, several limitations remain. First, our study examined a fixed developmental time window. The responsiveness of retinal progenitors to light likely changes over time, and determining the optimal timing, frequency, and duration of stimulation will be critical for clinical and translational applications. Second, we used wild-type human pluripotent stem cells; it remains to be seen whether photostimulation can rescue defects in disease-specific models, including retinal dystrophies or ciliopathies.

Taken together, our findings demonstrate that gamma-frequency light flicker can be used as a novel, physiologically aligned stimulus to promote photoreceptor maturation. This approach improves both cellular structure and neural output in retinal organoids, offering new opportunities for disease modelling, drug screening, and future translational applications such as photoreceptor transplantation.

## Methods

### Pluripotent stem cells culture and retinal organoids generation

We used human induced pluripotent stem cells (hiPSCs; cell line M8^37–39^) and human embryonic stem cells (hESCs; cell line H9) for retinal organoid generation. hiPSCs were cultured in Essential 8 medium (Gibco) on Vitronectin-coated dishes and passaged approximately every 4–5 days. hESCs were maintained in mTeSR™1 medium (STEMCELL Technologies) on Matrigel-coated plates (Corning), with routine passaging at similar intervals. Retinal organoids were derived from both cell lines using previously established protocols^34,40^ (**Fig. S1a**).

### Photostimulation of retinal organoids

Retinal organoids between day 150 and day 180 of the differentiation process were used, unless stated otherwise. One day prior to photostimulation, organoids were transferred into a black 96-well plate with a clear, round, ultra-low attachment bottom (#4515, Corning), each well containing fresh culture medium without phenol red and a single organoid. Photostimulation was performed using the CellLighteR device^20^. The LED module (**Fig. 1c**) was placed inside a standard tissue culture incubator, and the 96-well plate was positioned directly on top of the module. A black lid was used to minimize ambient light interference. Photostimulation began the day after seeding. The light intensity used for the stimulation was 15 lux of white light.

### Retinal organoid dissociation for single cell RNA sequencing

Organoids were photostimulated under a 12-hour light/12-hour dark cycle for one week, with the light phase consisting of 40 Hz flickering illumination. On day 7, organoids were collected during the dark phase to avoid acute light-induced effects, and all subsequent steps were performed under light-protected conditions. For each condition, 30 organoids were processed for single-cell RNA sequencing. Organoid dissociation was carried out using the Neurosphere Dissociation Kit (#130-095-943, Miltenyi Biotec) according to the manufacturer’s instructions. Briefly, organoids were washed twice with cold HBSS (#14025092, Gibco) and transferred into Eppendorf tubes containing 960 μL of preheated Buffer X (each set of 15 organoids was dissociated in a separate tube). Then, 25 μL of Enzyme P, 10 μL of Buffer Y, and 5 μL of Enzyme A were added and mixed gently. The tubes were placed on a thermal mixer with a block (Eppendorf) for 10 minutes at 37°C and 800 rpm. The organoids were dissociated using a fire-polished Pasteur pipette by pipetting up and down 10 times slowly. They were then incubated again under the same conditions for 5 minutes in the thermal mixer and pipetted up and down 10 times with the Pasteur pipette. After adding cold HBSS containing 5% FBS to the single-cell suspension, the mixture was passed through a 40 μm cell strainer placed on a 15 mL tube. Cells were counted, and the tubes were centrifuged at 300×g for 10 minutes at room temperature. The supernatant was aspirated completely, and the cells were resuspended in cold HBSS containing 5% FBS at a concentration of 1000 cells/μL and kept on ice. (Total single cell number 1.500.000/sample)

### Single cell RNA sequencing; library preparation and data processing

The cell concentrations were adjusted to 1,000 cells/µL. These suspensions were loaded onto the Chromium Controller following the Chromium Next GEM Single Cell 3’ Gene Expression v3.1 (Dual Index) protocol, Rev E (CG000315) provided by 10x Genomics. Library preparation was conducted using the same protocol. The quality and quantity of the prepared libraries were assessed using the High Sensitivity NGS Fragment Analysis Kit (Agilent Technologies) and the QuantiFluor dsDNA System (Promega). The libraries were then sequenced using the AVITI sequencing platform (Element Biosciences), employing the AVITI 2×75 Sequencing Kit Cloudbreak FS High Output (150 Gb). Sequencing reads were processed using Cell Ranger software version 8.0.1 (10x Genomics), with FASTQ files aligned to the human reference genome refdata-gex-GRCh38-2020-A, also provided by 10x Genomics, to generate gene expression matrices for downstream analyses. All data analysis and visualizations were conducted using R. We performed quality control checks for each sample in R for each sample and removed any cells with <1000 reads or 500 genes or >10% mitochondrial reads. Doublets were predicted using DoubletFinder^41^ and filtered from the data. The Seurat R package (version 5.1.0)^42^ was then used to process the data. The following R packages were used to further analyse and visualize the data: SeuratExtend^43^, scCustomize, clusterProfiler^44^, monocle3^45^, and ggstatsplot^46^.

### RNA isolation

For RNA extraction, a minimum of six organoids per sample were collected and thoroughly washed with phosphate-buffered saline (PBS). Organoids were then homogenized in 300 µL of RNA Blue Reagent (#R013, Top-Bio), a Trizol analogue, using a 1 mL insulin syringe to ensure complete cell lysis. After homogenization, total RNA was extracted using the Direct-zol™ RNA Microprep Kit (#R2062, Zymo Research) according to the manufacturer’s protocol.

### RT-qPCR analysis

For gene expression analysis, RNA was reverse transcribed into complementary DNA (cDNA) using the High-Capacity cDNA Reverse Transcription Kit (#4368813, Applied Biosystems), following the manufacturer’s instructions. Reverse transcription was carried out in a 20 µL reaction volume with incubation steps at 16 °C for 30 minutes, 42 °C for 30 minutes, and 85 °C for 5 minutes. The resulting cDNA was amplified using the LightCycler 480 Real-Time PCR System (Roche) and PowerUp SYBR Green Master Mix (#P552, Top-Bio) in 20 µL reactions. PCR cycling conditions included an initial denaturation at 95 °C for 10 minutes, followed by 40 cycles of 95 °C for 15 seconds and 60 °C for 1 minute. Primer sequences are listed in **Table S2**. Gene expression levels were normalized to *HPRT1* mRNA as the endogenous control. All experiments included at least three independent biological replicates, each performed in technical triplicate. Individual data points are shown in each graph. Statistical significance was assessed using a two-tailed Student’s t-test, with p-values indicated in the corresponding figure panels. Comparisons with p-values < 0.05 were considered statistically significant.

### Bulk RNA sequencing

RNA integrity was assessed using the Fragment Analyzer with the RNA Kit 15 nt (Agilent Technologies). For library preparation, 500 ng of total RNA was used as input for the QuantSeq 3′ mRNA-Seq FWD with UDI 12 nt Kit (v2) (Lexogen), in combination with the UMI Second Strand Synthesis Module for QuantSeq FWD. Library quality and quantity were evaluated using the QuantiFluor dsDNA System (Promega) and the High Sensitivity NGS Fragment Analysis Kit (Agilent Technologies). The final pooled libraries were sequenced on an Illumina NextSeq platform using the High Output Kit v2.5 (75 cycles) in single-end mode. Raw single-end FASTQ reads were quality-checked using FastQC. Adapter removal and quality trimming were performed with Trimmomatic v0.39 using the following parameters: CROP:250, LEADING:3, TRAILING:3, SLIDINGWINDOW:4:5, and MINLEN:35. Trimmed reads were aligned to the human reference genome (Ensembl GRCh38 annotation) using STAR v2.7.3a with default settings, except for -- outFilterMismatchNoverLmax 0.66 and -- twopassMode Basic. Post-alignment quality control—including assessment of uniquely and multi-mapped reads, rRNA contamination, mapping distribution, read coverage, strand specificity, gene biotypes, and PCR duplication— was performed using RSeQC v4.0.0, Picard toolkit v2.25.6, and Qualimap v2.2.2. Downstream analysis of the NGS datasets was conducted in RStudio using the following R packages: DESeq2, biomaRt, Rsubread, and clusterProfiler.

### Immunohistochemistry

Organoids were washed with PBS and fixed in 4% paraformaldehyde (PFA) solution in PBS for 30 minutes at room temperature. After fixation, the PFA was removed, and organoids were washed three times with PBS. They were then incubated overnight in 30% sucrose (Sigma-Aldrich) solution at 4 °C for cryoprotection. Subsequently, organoids were transferred into plastic disposable base molds, the remaining sucrose solution was carefully removed, and the molds were filled with Tissue-Tek® O.C.T. Compound medium (Sakura). The molds containing organoids were placed on dry ice for 15 minutes to solidify the medium and stored at −20 °C or processed immediately. Frozen blocks were sectioned into 7 µm-thick slices using a Leica CM1850 Cryostat (Leica). Slides were allowed to dry at room temperature for 20 minutes before further processing. For immunofluorescent staining, sections were washed with PBS and blocked for 1 hour at room temperature in a humidified chamber using a blocking buffer containing 0.3% Triton X-100 and 5% normal goat serum in PBS. Primary antibodies are shown in **Table S3** diluted in antibody diluent (AD, PBS with 0.3% Triton X-100 and 1% bovine serum albumin) were applied overnight at 4°C. After washing with AD, secondary antibodies (Alexa Fluor 488 or 594-conjugated Goat anti-Mouse/Rabbit IgG, Invitrogen, 1:1000) were applied for 1 hour at room temperature in a humidified chamber. Nuclei were stained with DAPI (1 µg/mL in PBS) for 5 minutes, and sections were mounted using Fluoromount Aqueous Mounting Medium (#F4680, Sigma Aldrich). Outer segment length was measured using ImageJ software, defined as the distance from the edge of the DAPI signal to the distal end of the Arrestin-C and ROM1 signals.

### Scanning electron microscopy

Retinal organoids were initially fixed in 3% glutaraldehyde (#G5882, Sigma-Aldrich) prepared in 0.1 M cacodylate buffer (pH 7.3; #C0250, Sigma-Aldrich). Following fixation, organoids were rinsed with fresh 0.1 M cacodylate buffer to remove residual fixative. Dehydration was performed through a graded ethanol series (30%, 50%, 70%, 80%, 90%, 96%, and absolute ethanol). After dehydration, organoids were dried using a critical point dryer (CPD 030, BAL-TEC Inc.). Dried organoids were then coated with a thin layer of gold using a Bal-Tec SCD 040 sputter coater (LEICA). The gold-coated samples were imaged using a scanning electron microscope (VEGA TS 5136 XM, Tescan Orsay Holding).

### Transmission electron microscopy

Organoids were fixed in 3% glutaraldehyde (#G5882, Sigma-Aldrich) in 0.1 M cacodylate buffer (pH 7.3; #C0250, Sigma-Aldrich) for 24 hours at 4 °C. After fixation, samples were washed with 0.1 M cacodylate buffer and post-fixed in 1% OsO_4_ (#05500, Sigma-Aldrich) in the same buffer for 1.5 hours at room temperature. Following post-fixation, samples were dehydrated through a graded ethanol series (50%, 70%, 96%, and absolute ethanol) and embedded in Durcupan™ ACM resin (Sigma-Aldrich). Polymerization was carried out over 3 days with a gradual temperature increase from 60 °C to 80 °C. Resin blocks were processed according to standard protocols for transmission electron microscopy, sections were prepared on Leica EM UC6 (Leica). Ultrathin sections were examined using a Morgagni 268D transmission electron microscope (Thermo Fisher Scientific) operating at 90 kV and equipped with a Veleta CCD camera (Olympus).

### Western blot analysis

For each sample, at least six organoids were rinsed three times with PBS and lysed in a buffer containing 50 mM Tris-HCl (pH 6.8), 10% glycerol, and 1% SDS. Lysates were sonicated to ensure complete homogenization, and protein concentrations were determined using the DC Protein Assay (Bio-Rad). Subsequently, lysates were mixed with 0.01% bromophenol blue and 1% β-mercaptoethanol, then heated at 100 °C for 5 minutes to denature proteins. Equal amounts of protein were loaded onto SDS-polyacrylamide gels for electrophoresis, followed by transfer to PVDF membranes (Merck Millipore) pre-activated with methanol. Membranes were blocked for 1 hour at room temperature in 5% skimmed milk prepared in TBST. Primary antibodies (**Table S3**), diluted in either 5% skimmed milk or 5% BSA (Sigma-Aldrich) in TBST, were applied overnight at 4 °C. The following day, membranes were washed three times for 15 minutes with TBST and incubated for 1 hour at room temperature with the appropriate secondary antibodies (**Table S3**). Protein bands were visualized using the ECL Plus Reagent Kit (GE Healthcare).

### Electrophysiological recordings

Electrophysiological recordings were performed as described previously^20,21^. The organoids were placed in artificial cerebrospinal fluid (aCSF), containing: 118 mM NaCl, 25 mM NaHCO_3_, 1 mM NaH_2_PO_4_, 3 mM KCl, 1 mM MgCl_2_, 2 mM CaCl_2_, 10 mM glucose, and 0.5 mM L-Glutamine. The aCSF was equilibrated with 95% O2 and 5% CO2. The organoids were then longitudinally opened and placed with the presumed RGC layer facing down onto a 4096 channel multielectrode array (MEA). A translucent polyester membrane filter (Sterlitech Corporation) was used to flatten the organoids. The organoids were allowed to settle for a minimum of 1 hour. Recordings were performed on the BioCam4096 MEA platform with BioChips 4096S+ (3Brain). The platform contained 4096 square microelectrodes in a 64 × 64 array configuration. Spike extraction was done using a quantile-based event detection method^47^. An automated spike sorting method for dense, large-scale recordings was used to sort single-unit spikes^48^. Statistical analysis and firing rate analysis were conducted using Prism (GraphPad) and MATLAB (MathWorks). cGMP (8-Bromoguanosine 3′,5′-cyclic monophosphate, # B1381, Sigma-Aldrich) was puffed into the recording chamber at a final concentration of 100 μM. Activity was recorded continuously for 10 minutes, starting with the spontaneous activity recording (baseline, 5 min) followed by the cGMP puff recording (5 min).

### Circadian rhythms measurement

For BMAIL1 expression monitoring using luciferase reporters, hiPSCs were transduced using lentiviral particles containing the pABpuro-BluF vector (pABpuro-BluF was a kind gift from Steven Brown [plasmid #46824; Addgene; http://n2t.net/addgene:46824; RRID:Addgene_46824])^49^. Lentiviral particles were generated, as described in^38,50,51^. Upon transduction, hiPSCs BMAL1::Luc were cultured in the presence of Puromycin (0.5 μg/ml).

For dexamethasone synchronization, the culture medium was replaced with pre-warmed medium containing 100 nM dexamethasone (#D4902, Merck). Organoids were incubated at 37 °C for one hour to allow synchronization. Following incubation, the dexamethasone-containing medium was removed, and the cells were washed three times with warm PBS to eliminate any residual compound. The PBS was then replaced with fresh, pre-warmed culture medium.

For luciferase activity measurement we used a modified version of LuminoCell^25^. This version is capable to measure luciferase activity in individual wells of 96-well plate (**Fig. 5a**). Individual organoids were seeded into wells of a black 96-well plate with a clear, round, ultra-low attachment bottom (#4515, Corning) in the presence of 100 μM D-Luciferin sodium salt (#L6882; Merck). The LuminoCell was placed into a tissue incubator, the 96 well plate was positioned on the luminometer and covered by a lid to protect the cells from potential incoming light. Luciferase activity was measured using a built-in serial monitor in the Arduino IDE software on a laptop computer Lenovo SL500 with an Ubuntu operating system (20.04 version). The measured values were saved into a CSV file and processed in R studio.

## Data Availability

NGS data has been deposited to GEO.

https://www.ncbi.nlm.nih.gov/geo/query/acc.cgi?acc=GSE299921

## Acknowledgements

This publication was supported by the Czech Science Foundation (GA21-08182S), by the Ministry of Health of the Czech Republic, grant nr. NU22-07-00380, by the Faculty of Medicine MU (MUNI/A/1738/2024), and by the European Union (Grant Agreement No. 101059788). We acknowledge the core facility CELLIM supported by MEYS CR (LM2023050 Czech-BioImaging). We acknowledge the CF Genomics and the CF Bioinformatics supported by the NCMG research infrastructure (LM2023067 funded by MEYS CR) for their support with obtaining scientific data presented in this paper. M.L. would like to acknowledge funding from the MRC UK (MR/X001687/1) and EPSRC UK (EP/Y031016/1)

We are deeply grateful to Professor Evelyne Sernagor for her invaluable assistance with electrophysiological measurements and for her insightful guidance throughout this project. Her pioneering contributions to the field of retinal neuroscience have left a lasting legacy. We are honoured to have had the opportunity to collaborate with her. Evelyne’s recent passing is a profound loss to the scientific community, and she will be greatly missed.

## Author Contributions

C.C. and T.B. conceived and designed the study; C.C., E.H., K.K., B.H., F.M.G., K.W., L.E., N.V., M.S., performed experiments and analysed data, under the supervision of T.B., M.L. and E.S.; T.B. analysed the scRNA-seq data; B.H., M.S., G.H., E.S. designed and performed the electrophysiological experiments, with G.H. analysing the data; C.C, G.H., E.S., M.L., and T.B. interpreted data; T.B. and C.C. wrote the manuscript, with feedback from M.L. and A.G.; All authors reviewed and edited the manuscript.

## Competing interests

The authors declare no competing interests.

## Supplementary Figures

**Figure S1.**
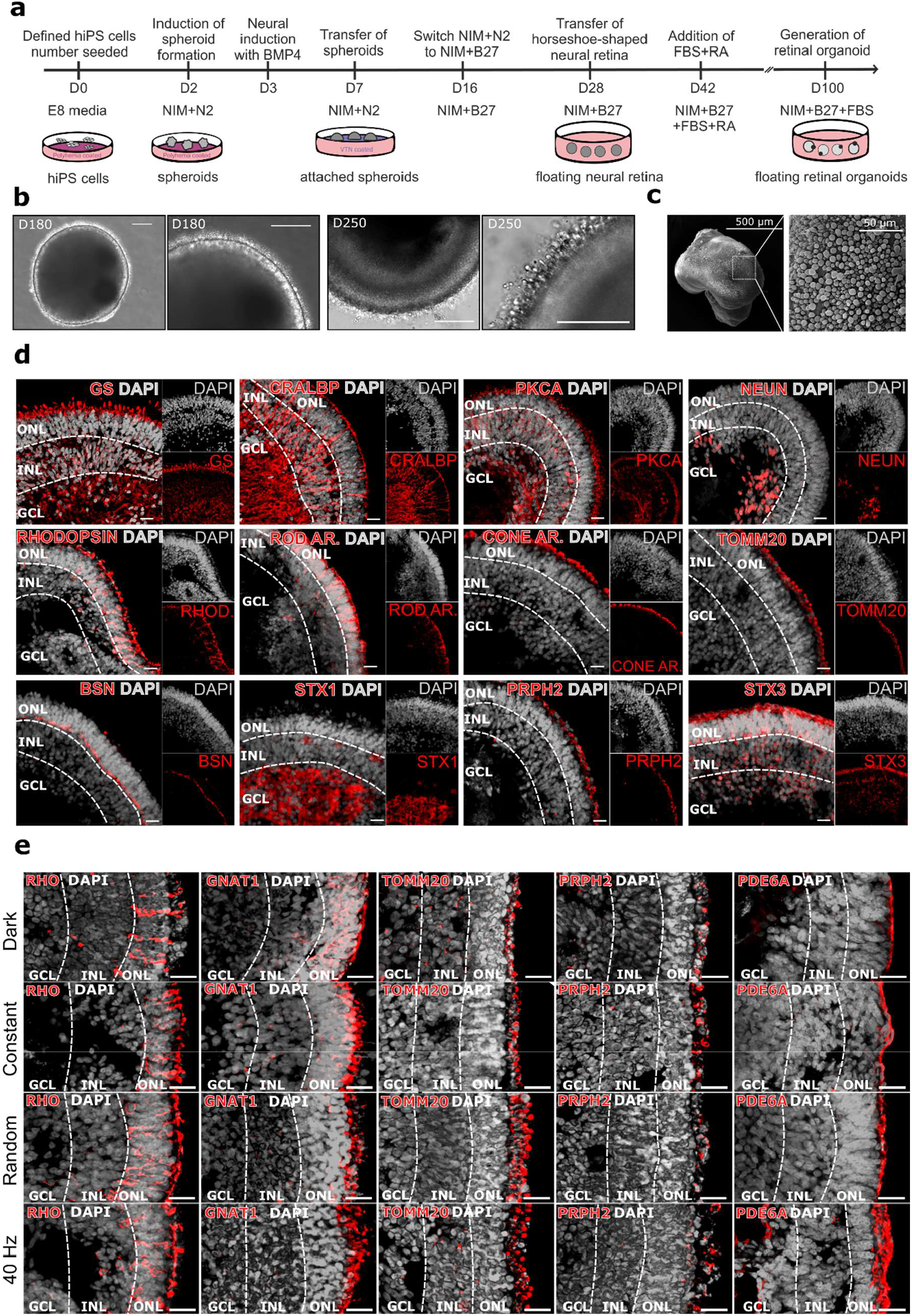
Characterization of human retinal organoids. To establish a robust foundation for our study, we first evaluated the structural integrity of our human retinal organoids. The results demonstrate that our differentiation protocol successfully generates highly organized and mature retinal organoids, closely recapitulating the architecture of the human retina. **(a)** The schematic overview of the differentiation protocol of hiPSCs. **(b)** Brightfield imaging reveals the characteristic morphology of our organoids, including well-defined brush borders, a hallmark of developing outer segment (OS)-like structures, indicating that photoreceptors are present and are also undergoing the expected morphological maturation required for functional light sensitivity. Scale bar = 200 μm **(c)** Scanning electron microscopy (SEM) confirms the presence of densely packed OS-like protrusions. **(d)** Immunostaining further verifies the presence of major retinal cell types and specialized substructures, including: glutamine synthetase (GS), with rods specifically marked by RHODOPSIN and ROD ARRESTIN (ROD AR.), and cones by CONE ARRESTIN (CONE AR.). Müller glial cells are labelled with CRALBP, while bipolar cells express PKCα, and retinal ganglion cells (RGCs) are marked by NEUN, confirming the presence of a complete neuronal network. Further, TOMM20 labelling highlights inner segment (IS) structures, and peripherin-2 (PRPH2) marks rod and cone outer segment organization, further demonstrating advanced photoreceptor differentiation. Finally, the expression of BSN and STX1 in ribbon synapses, as well as STX3 in conventional synapses, indicates the formation of functional synaptic connections, supporting the notion that these organoids are not only structurally developed but also capable of neural communication. Scale bars = 20 μm. ONL = outer nuclear layer, INL = inner nuclear layer, GCL = ganglion cell layer. **(e)** Representative immunostaining images of RHODOPSIN (RHO), GNAT1, TOMM20, PRPH2, and PDE6A under all tested photostimulation regimes. Scale bar = 20 μm. ONL = outer nuclear layer, INL = inner nuclear layer, GCL = ganglion cell layer.

**Figure S2:**
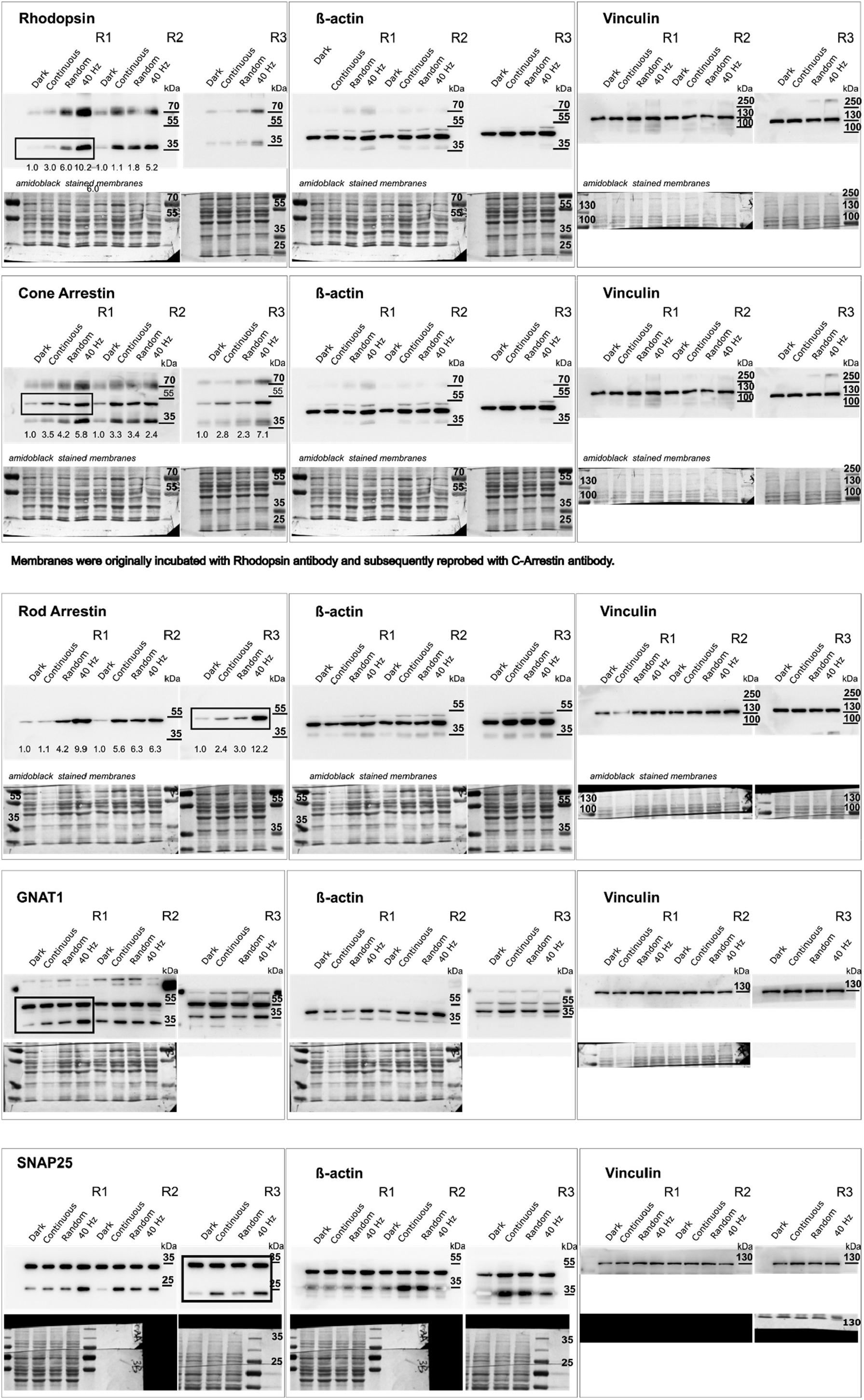
Western Blot full scans. R1, R2, and R3 represent individual replicates.

**Figure S3:**
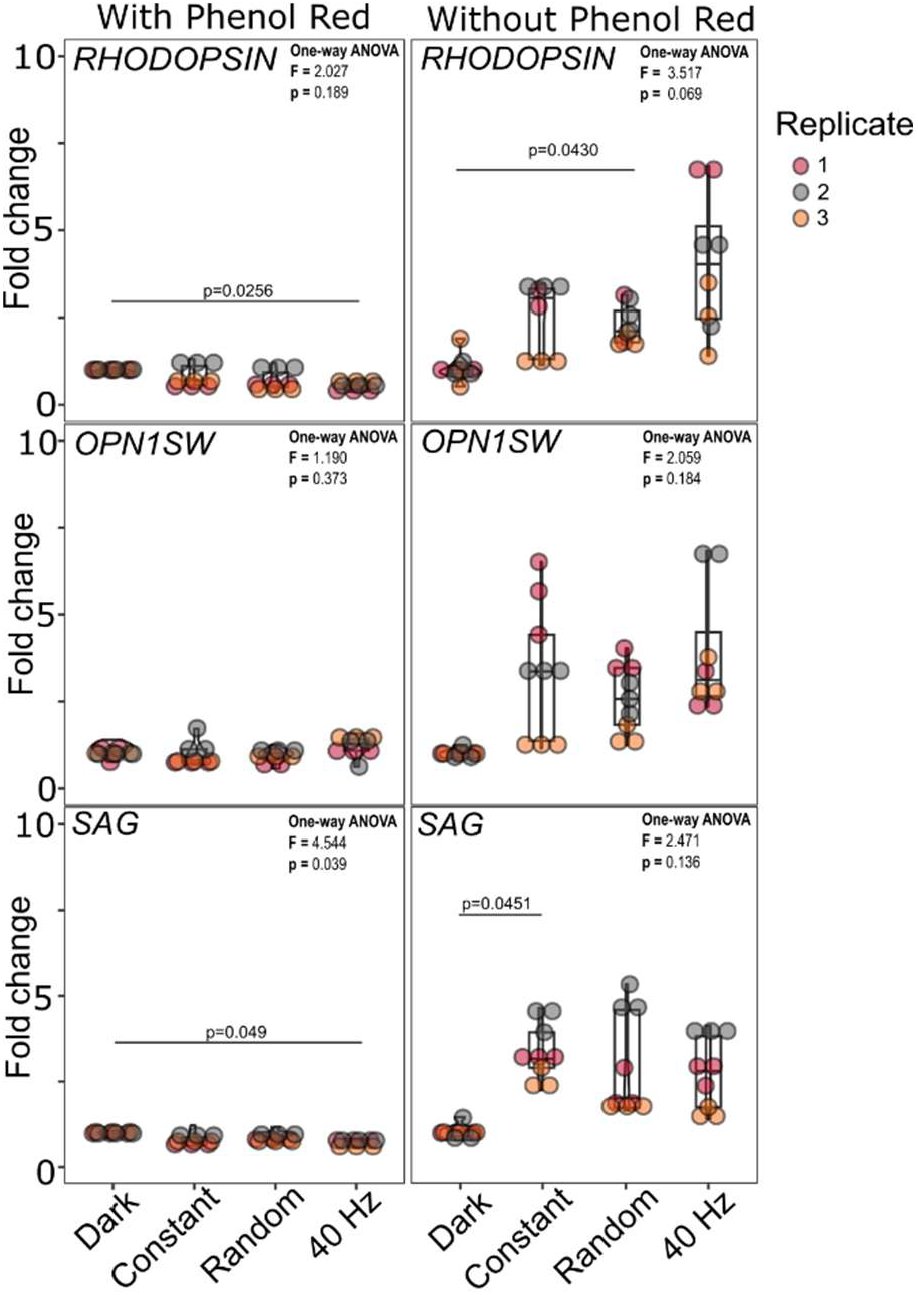
The presence of phenol red in culture medium diminishes the effect of photostimulation on the expression of OS-specific genes. Graphs shows expression of RHODOPSIN (RHO), OPN1SW, and SAG, as demonstrated by RT-qPCR. Statistics – one-way ANOVA, post hoc t-test.

**Figure S4:**
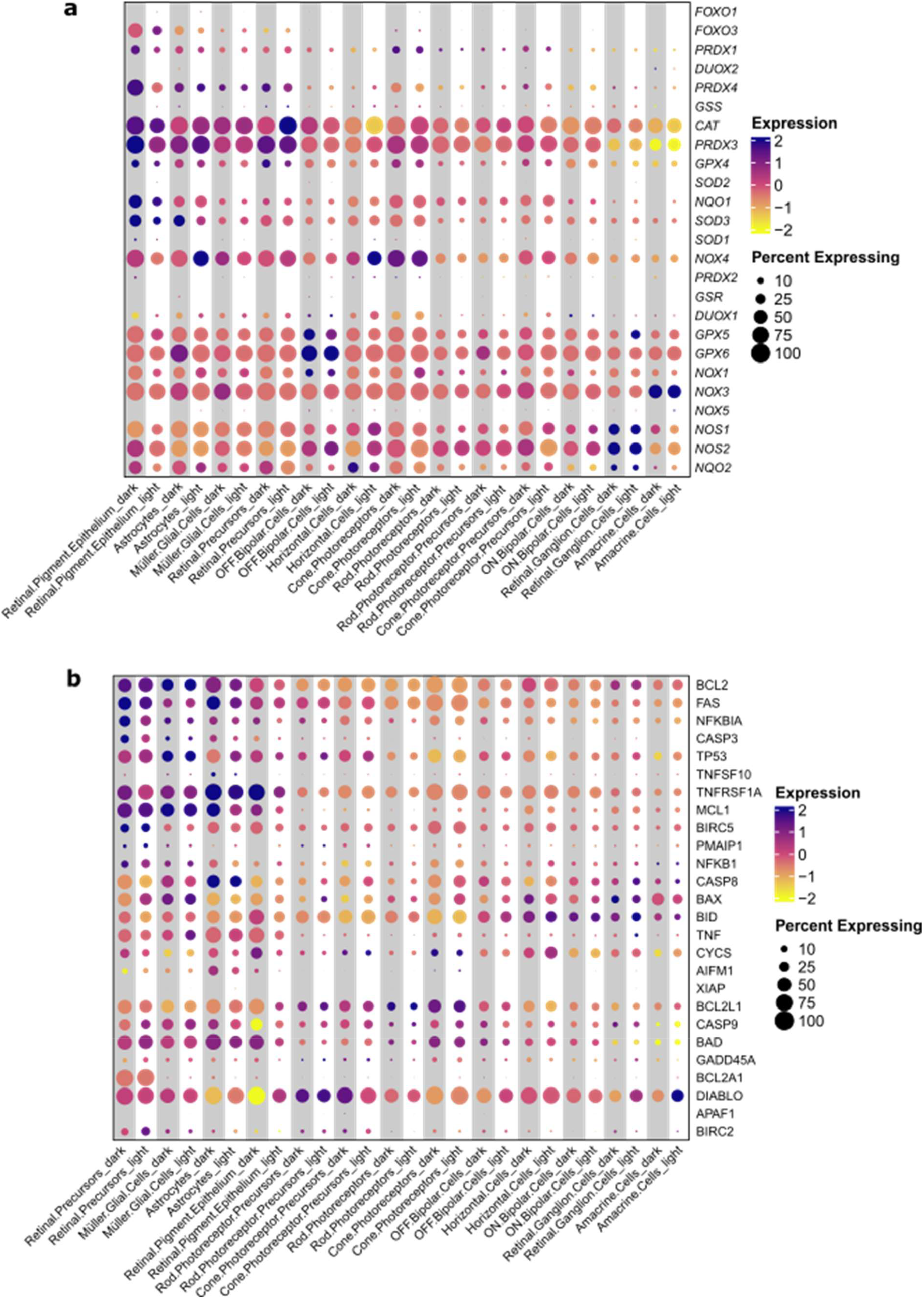
Photostimulation does not upregulate the expression of major genes involved in oxidative stress and apoptosis. Dot plots showing expression of major genes in **(a)** oxidative stress and **(b)** apoptosis upon photostimulation of retinal organoids.

**Figure S5.**
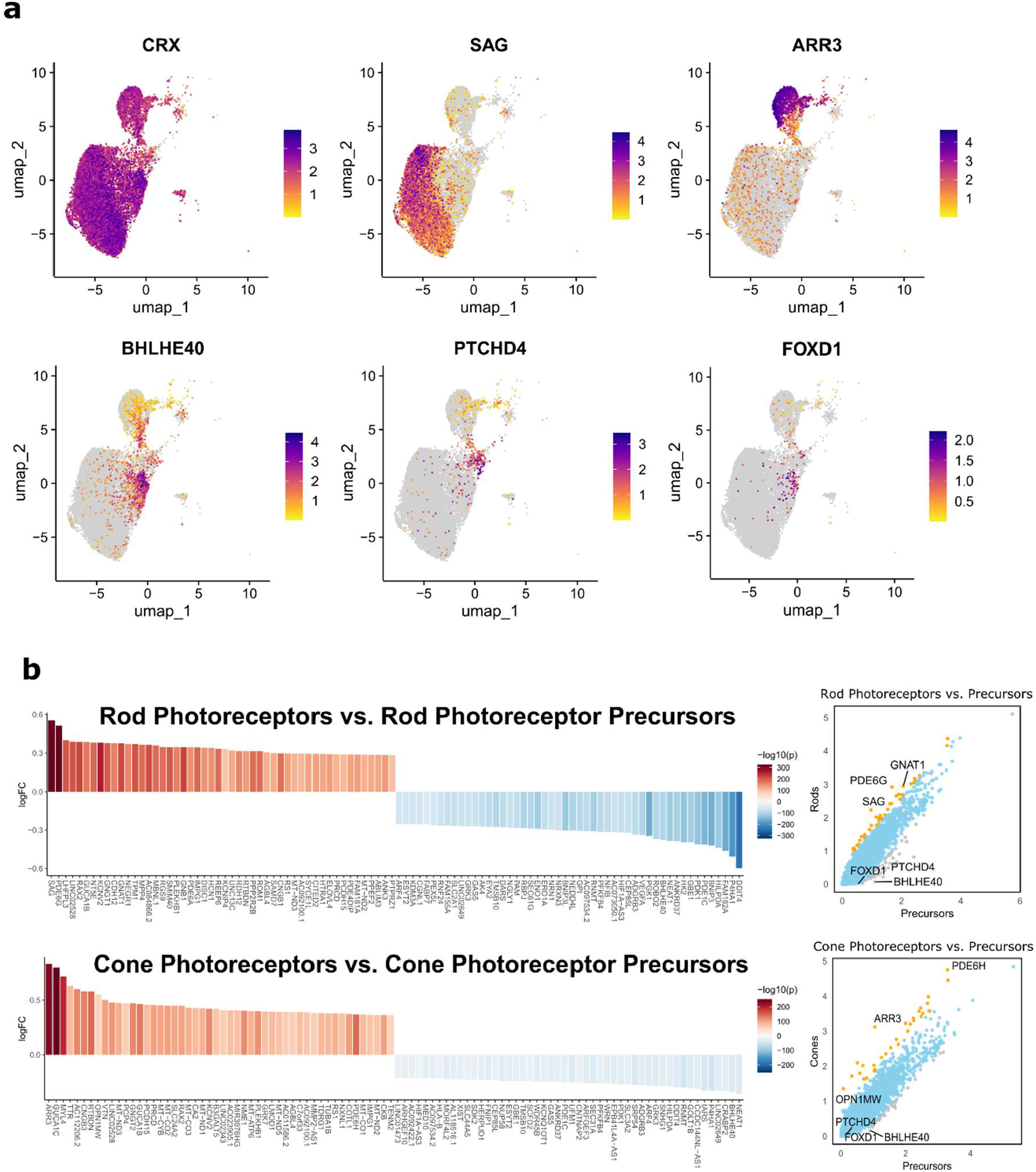
Gene expression signatures distinguish mature photoreceptors from their precursors. **(a)** UMAP feature plots showing expression of key marker genes across the photoreceptor population. CRX is broadly expressed in both precursor and mature photoreceptors. In contrast, SAG and ARR3 mark mature rod and cone photoreceptors, respectively, while BHLHE40, PTCHD4, and FOXD1 are selectively enriched in precursor populations. **(b)** Waterfall plots display the top 50 upregulated and top 50 downregulated genes in mature rods versus rod precursors and mature cones versus cone precursors, highlighting major transcriptional differences associated with photoreceptor maturation. Adjacent dot plots further visualize differential gene expression patterns between mature and precursor states.

**Figure S6.**
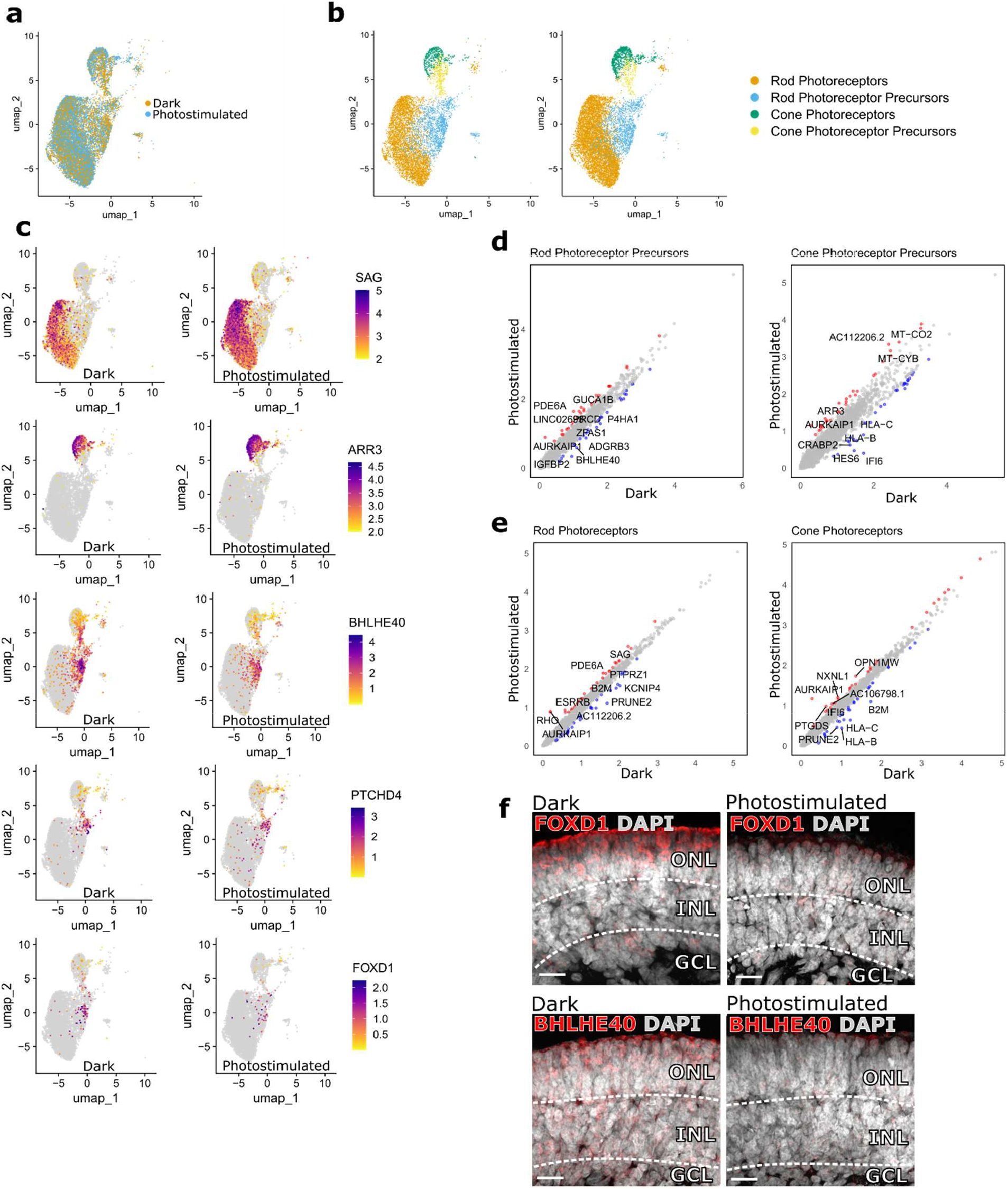
Photostimulation enhances photoreceptor maturation and alters subtype-specific gene expression. **(a)** UMAP embedding of all photoreceptors with overlay of cell identities by condition: yellow dots indicate cells from dark-cultured organoids; green dots represent photostimulated cells. **(b)** Separate UMAP plots of photoreceptors from dark (left) and photostimulated (right) conditions, showing the distribution across four clusters: rod photoreceptors, cone photoreceptors, rod precursors, and cone precursors. Photostimulation increases the representation of mature photoreceptor clusters. **(c)** UMAP feature plots displaying gene expression of SAG, ARR3, BHLHE40, PTCHD4, and FOXD1 in dark (left) and photostimulated (right) conditions. Mature markers are upregulated in stimulated cells, while precursor markers are downregulated. **(d)** Dot plots showing gene expression in rod precursors (left) and cone precursors (right) under dark (x-axis) and photostimulated (y-axis) conditions. **(e)** Equivalent plots for mature rods (left) and cones (right), highlighting robust upregulation of photoreceptor-specific genes upon photostimulation. **(f)** Representative immunostaining images of BHLHE40 and FOXD1 under dark and photostimulation regimes. Scale bar = 20 μm. ONL = outer nuclear layer, INL = inner nuclear layer, GCL = ganglion cell layer.

**Figure S7.**
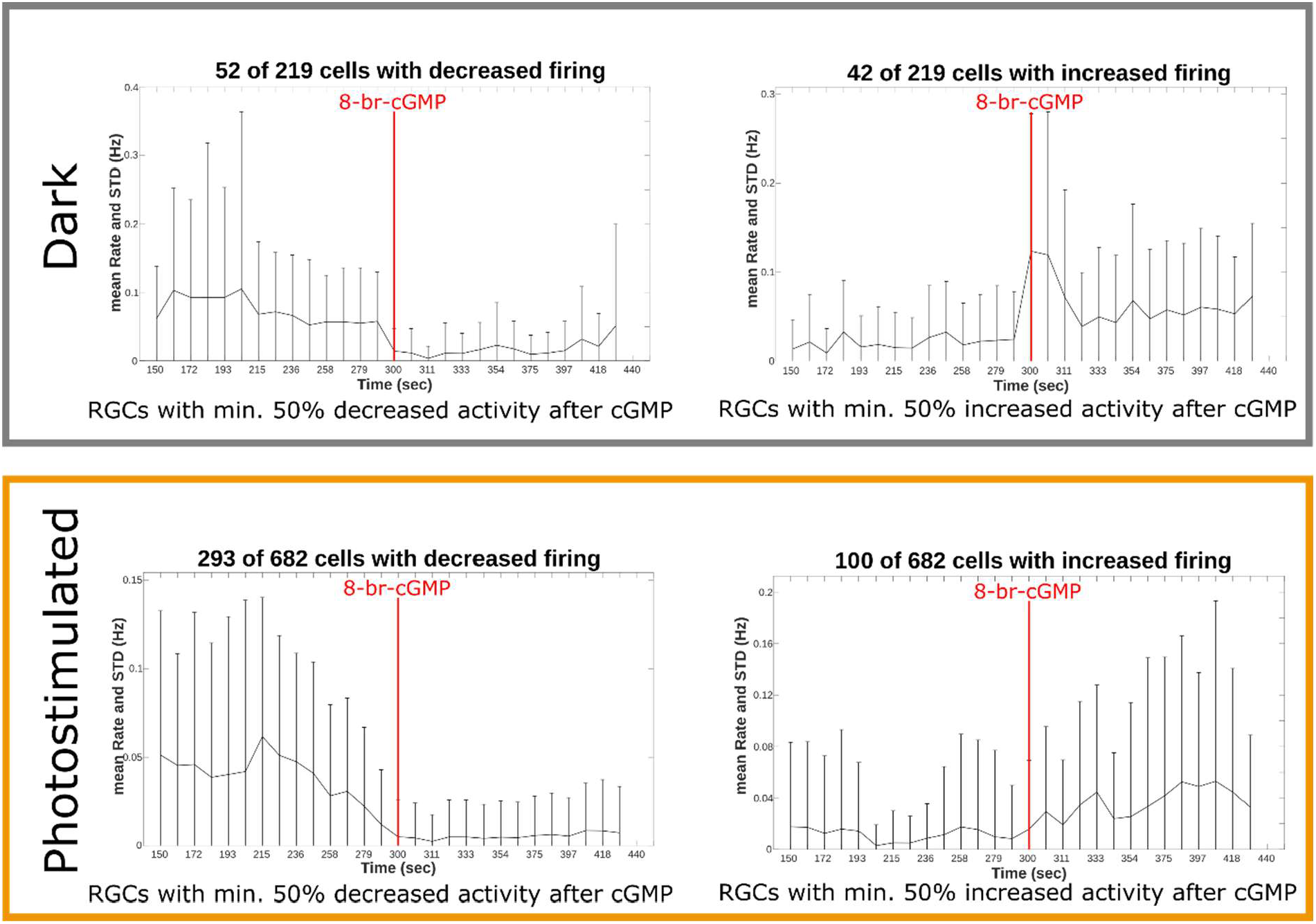
Photostimulation enhances RGC responses. Average firing rates of RGCs over time before and after 8-br-cGMP application in dark (top) and photostimulated (bottom) organoids. Only RGCs exhibiting a ≥50% change in activity (either increase or decrease) following stimulation were included. Time of 8-br-cGMP addition is shown as a red bar in the graph.

## Notes

https://www.ncbi.nlm.nih.gov/geo/query/acc.cgi?acc=GSE299921

### Competing Interest Statement

The authors have declared no competing interest.

